# Phylogenetic discordance can substantially overestimate genomic reassortment in avian influenza virus

**DOI:** 10.1101/2025.02.24.639875

**Authors:** Hugo G. Castelán-Sánchez, Art F. Y. Poon

**Affiliations:** Department of Pathology & Laboratory Medicine, Western University, Dental Sciences Building, Rm. 201, London, Ontario N6A 5C1, Canada; Department of Microbiology & Immunology, Western University, 1151 Richmond Street, London, Ontario N6A 3K7, Canada; Department of Computer Science, Western University, Rm. 355, Middlesex College, London N6A 5B7, Canada

## Abstract

Recombination plays an important role in the evolution of RNA viruses, as it allows the exchange of genetic material between viral lineages. Reassortment, a form of recombination specific to segmented genomes, involves the exchange of entire segments and has contributed significantly to the adaptation and spread of influenza viruses through novel genomic combinations, *i.e*., antigenic shifts. It is usually identified by phylogenetic discordance: differences in the topologies of trees reconstructed from different genomic segments. However, phylogenetic discordance can also result from error in reconstructing trees. To characterize the impact of reconstruction error, we curated a database of *n* = 11, 765 complete genomes of avian H5Nx influenza A viruses from avian hosts. We found evidence of widespread reassortment as measured by inferred subtree-prune-regraft (SPR) events, consistent with previous studies. Next, we ran replicate simulations of sequence evolution along the reference tree for the segment encoding hemagglutinin (HA), adjusting simulations for the lengths and clock rates of the other segments. These simulations provided a baseline for the expected amount of phylogenetic discordance in the absence of any reassortment. When sampling HA sequences at random from the database to build reference trees, we observed that simulating other segments without reassortment still yielded about 32% as many SPRs as the real segment data on average. The average proportion of SPRs without reassortment was greatly reduced (4%) if we selected an equivalent number of HA sequences retaining the most genetic diversity, which was consistent with the accuracy of phylogenetic reconstruction being the limiting factor. This implies that measuring reassortment by SPRs may have a high false positive rate, and that previous evidence of extensive reassortment in influenza viruses should be interpreted with caution. In addition, we observed that the SPRs reconstructed on simulated trees had significantly shorter distances between the prune and regraft locations than real trees. These results suggest that down-sampling sequences to maximize evolutionary divergence and filtering out the shortest SPRs may be effective measures against false positives.

## Introduction

Recombination is an important mechanism of genomic evolution in RNA viruses (Simon-Loriere and Holmes 2011). This process involves the exchange of genetic material between different genomic backgrounds, which is facilitated by the coinfection of a host by two or more distinct virus lineages. Recombination uncouples the evolutionary fate of a mutation from the rest of the genome, and under certain conditions (Otto and Barton 1997), recombination will increase the diversity of genotypes in the population (Pérez-Losada et al. 2015; Wang et al. 2022). Many RNA viruses have segmented genomes, including members of the *Reoviridae* and *Orthomyxoviridae* families (McDonald et al. 2016). In these viruses, recombination can occur by reassortment, in which the genomic segments packaged into a virion are derived from different parental genomes. Additionally, many other viruses have multipartite genomes where segments are packaged into separate virus particles. These viruses can reassort segments from different infected cells or even from different hosts (Varsani et al. 2018). Reassortment involves the wholesale exchange of entire segments between genomes, which can accelerate the acquisition of combinations of mutations that are required to overcome host barriers more rapidly than the progressive accumulation of individual mutations (Vijaykrishna et al. 2015).

Influenza A virus (IAV; family *Orthomyxoviridae*, genus *Alphainfluenza*) provides several important examples of the possible impacts of genetic reassortment on the health of human and non-human animal populations. This virus has been responsible for numerous and severe epidemics and pandemics (Sutton 2018), several of which are associated with a reassortant virus (Lowen 2017). IAV has a negative-sense RNA genome that consists of eight genomic segments that encode 10 major proteins and other accessory gene products (Pinto et al. 2021). This number of segments implies that about 99% virus particles produced by a co-infected host cell will contain a reassortant genome. The co-circulation of multiple IAV subtypes and clades in domesticated animal host species provides abundant opportunities for reassortment between divergent lineages (Ryt-Hansen et al. 2021; Li et al. 2024). Thus, reassortment can generate new chimeric IAV genomes that potentially increase the capacity of this virus to be transmitted to a new host species and establish a productive infection (Bouvier and Palese 2008; Ma et al. 2016).

The comparative analysis of IAV genome sequences plays an important role in measuring the frequency of reassortment and detecting the emergence of new reassortant viruses (Kong et al. 2015; Taylor et al. 2023). To this end, a number of different methods have been developed to identify reassortment events. In the absence of recombination, the evolutionary history of a sample population of genome sequences may be represented by a single phylogenetic tree. Reassortment uncouples the evolutionary histories of the affected genomic segments. Consequently, a common technique to detect reassortment is to reconstruct a phylogeny for each segment, and then to identify discrepancies in the topology and/or labels among the resulting trees. Discordant trees are often visualized as tanglegrams (Page 2003), in which two trees with a common set of labels are drawn in opposing orientations, with subtrees rotated to maximize the concordance of tip labels between trees. Tanglegrams can be used to visually assess a small number of reassortment events. In addition, the optimization algorithm for untangling pairs of trees provides a rough measure of the extent of discordance that may be due to recombination, *i.e*., entanglement. However, tanglegrams do not readily scale to resolving large numbers of reassortment events. Evolutionary events that produce discordant trees can be explicitly modeled using an ancestral recombination graph (ARG; Wong et al. 2024). Although there has been a recent resurgence of interest in ARGs (Frost et al. 2015), they remain too computationally complex for large-scale application.

A compromise between these two extremes is to infer the frequency of reassortment events by the number of tree rearrangement operations required to convert one segment tree to another (Svinti et al. 2013), which can be interpreted as an edit distance. Screening for tips that transition between clusters of labels in the respective trees is an equivalent approach Silva et al. 2012; Gong et al. 2021. There are many edit distances available for this purpose, including the Robinson-Foulds distance (RF; Briand et al. 2020), tree bisection and reconnection (TBR; Kelk and Linz 2020), nearest neighbor interchange (NNI; Collienne and Gavryushkin 2021), and subtree prune and regraft (SPR; Wu 2009). Among these, the SPR distance is frequently used in studies of lateral gene transfer and recombination (Whidden et al. 2014), possibly because the SPR most closely resembles the effect of recombination on the topologies of trees on either side of the breakpoint.

In this work, we performed a phylogenetic analysis starting from nearly twelve thousand full genome sequences for influenza A H5Nx viruses to identify putative rearrangement events by reconstructing SPR rearrangements. Our main objective was to evaluate the reliability of this approach given the uncertainty in reconstructing accurate phylogenies under realistic conditions. To this end, we reconstructed phylogenies for each IAV H5Nx segment and then selected one of these trees as the template on which we simulated sequence evolution for the other segments in the absence of reassortment. We compared the entanglement values and the SPR distances between the reference tree for segment 4 (encoding hemagglutinin, HA) and trees for all other segments for both real and simulated data sets to assess the specificity of these methods for detecting reassortment. In addition, we characterized the SPR operations reconstructed from these data in terms of the size of the subtree and the distance traversed in the tree.

## Methods

### Data collection

We retrieved influenza A virus H5Nx genome sequences isolated from avian hosts from the NCBI Genbank and GISAID databases (accessed 5 May 2024). The GISAID query returned 5,994 to 6,000 sequences per segment, and the Genbank query returned 6,157 to 6,170 sequences per segment. We combined these sequence data sets by genomic segment. There was only a small proportion of sequences that were exact duplicates between databases. For example, out of 12,167 neuraminidase (NA) sequences, there were 8,347 unique sequences, of which 1,417 (17%) had two or more instances — only 171 (12%) of these were represented in both databases, involving a total of 673 sequence records. This low amount of overlap is largely due to a transition away from depositing sequences in Genbank to GISAID around 2020, coinciding with recent outbreaks of avian H5N1 IAV (Supplementary Figure S1). Sequence identity was not sufficient to determine whether the sequences are derived from the same isolate. Consequently, we merged these datasets by harmonizing the sequence labels into the World Health Organization (1980) nomenclature format, including host species, country of sampling, isolate name and year of sampling, and then filtering sequences with duplicate labels. This reduced the combined data to 11,795 sequences per segment. We generated alignments for each segment using MAFFT (v7.490; Katoh and Standley 2013), removed duplicate sequences, and then manually inspected and cleaned each segment alignment with trimAl (Capella-Gutiérrez et al. 2009, p. v1.5.0).

We reconstructed maximum likelihood phylogenies using the standard generalized time-reversible (GTR) model with gamma-distributed rate variation and empirical base frequencies IQ-TREE2 (Minh et al. 2020). To reduce the number of genomes, we applied two subsampling strategies to obtain subsets of sequences. First, we tested the prune function of the Biopython Phylo module to progressively remove terminal branches in the HA tree whose length exceeded a cutoff. After experimenting with different thresholds, we selected a cutoff of 0.08 expected substitutions per site, resulting in a subset of 213 HA sequences. In addition, we generated five independent random samples of 213 sequences per segment from the entire dataset. We used the labels of the sampled HA sequences to retrieve the corresponding sequences for the other seven segments and reconstruct trees. These two sampling strategies provided a means of evaluating the effect of phylogenetic reconstruction error on levels of phylogenetic discordance.

### Data simulation

To quantify the discordance among phylogenies reconstructed from sequences that have evolved in the complete absence of reassortment, we simulated sequence alignments along the tree inferred from the real HA alignment (*T*_HA_) for every other segment. Since every segment would evolve along exactly the same tree topology, any reassortment event inferred for the simulated trees could be interpreted as a false positive. We proportionally rescaled branch lengths in *T*_HA_ so that the total tree length corresponded to the length of the tree reconstructed from actual sequences of that segment. Our motivation for adjusting tree lengths in this manner was to accommodate the substantial variation in rates of evolution among segments, which could otherwise confound our assessment of false positive rates. For each rescaled tree, we used the Python module *Pyvolve* (Spielman and Wilke 2015) to simulate sequence evolution under the default Goldman-Yang codon substitution model with the dN/dS ratio set to 0.5 and sequence length matched to each genomic segment.

### Phylogenetic analysis

We used two approaches to infer reassortment events. First we used a tanglegram, which is a graphical comparison of two trees with a common set of tip labels. The trees are rotated and optionally re-rooted to minimize the discordance in the vertical location of labels, as quantified by the entanglement statistic. Our working assumption is that increasing amounts of reassortment will induce greater amounts of entanglement between trees of different segments. We used the *step2side* method implemented in the Python module *tanglegram* (Nguyen et al. 2022) to disentangle the HA tree and trees from every other segment. We further visualized the tanglegram using the cophylo function from the phytools package in R, which implements a closest-neighbor (NN) tanglegram approach by heuristically rotating the trees to minimize link crossings between matched tip labels (Revell 2024).

Second, we used the *Espalier* library (Rasmussen and Guo 2023) in Python to efficiently calculate the approximate number of SPR (subtree prune and regraft) operations required to convert one phylogenetic tree into another. This number provides another means of estimating the number of reassortment events between phylogenetic trees. We obtained SPR distances between the HA tree and the trees for every other segment for the actual IAV genome sequence data, and we made the same comparisons for trees reconstructed from simulated data.

We hypothesized that false positive reassortment events due to phylogenetic uncertainty would tend to involve SPR moves separated by shorter distances in the respective trees. To evaluate this hypothesis, we quantified the total path length separating the inferred graft points for each SPR operation. Let the tree reconstructed from real sequences for the *i*-th segment be denoted by *T*_*i*_. Moreover, let the tree reconstructed from simulated sequences that were evolved along the real HA tree *T*_4_ be denoted by 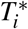. For each subtree *S* _*j*_ in the maximum agreement forest between trees *T*_4_ and *T*_*i*_ for *i* ≠ 4, we first locate *S* _*j*_ in the reference (segment 4) tree *T*_4_ by searching for the internal node with the fewest descendants that match all tip labels in *S* _*j*_. We call this node the root of subtree *j* in *T*_4_ and denote it as 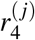. Next, we located the internal node in *T*_4_ with the same direct common ancestor as 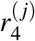, which we refer to as its sibling node, 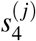. Switching to the other segment tree *T*_*i*_, we locate the same subtree *S* _*j*_ to identify 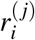 by matching tip labels. Finally, we locate the internal node in *T*_*i*_ with the fewest descendants that match all tip labels associated with 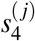, which we denote as 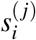. The sibling distance is then defined as the total path length (*i.e*., sum of branch lengths) from 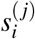 to 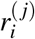. Put another way, 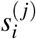 is the approximate location from which the SPR operation pruned the subtree, and 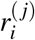 is the location where the subtree was grafted to. A diagram summarizing this procedure is provided as Supplementary Figure S2. Following this method, we calculated the sibling distances for all SPR events between *T*_4_ and both real (*T*_*i*_) and simulated 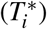 trees for other segments.

## Results

We obtained a total of 11,795 H5Nx influenza A virus genome sequences from the NCBI Genbank and GISAID databases. These sequences represented a variety of avian hosts, with the most frequent annotations being ‘chicken’ (*n* = 2, 968), ‘duck’ (*n* = 1, 774) and ‘turkey’ (*n* = 927). There was a total of 527 unique annotations for host organisms at varying taxonomic levels; for instance, mallard duck (*n* = 47) and Muscovy duck (*n* = 199) were both sub-classifications below ‘duck’. Similarly, there were 621 unique geolocation annotations, with the three most common being ‘Netherlands’ (*n* = 845), ‘Minnesota’ (*n* = 569), and ‘Vietnam’ (*n* = 557). Many records were labeled by state instead of country, obscuring a relative abundance of samples from China and the United States, for example. We reconstructed a maximum likelihood phylogeny relating all HA (segment 4) sequences, and then we generated a sub-sample of *n* = 213 (1.8%) sequences maximizing the total tree length by progressively removing the shortest terminal branches. Next, we extracted the corresponding sequences for the other segments (*i.e*., from the same virus isolates as the HA sample) and regenerated maximum likelihood phylogenies for each segment. These seven datasets are intended to minimize the error in phylogenetic reconstruction for a given sample size, and will be referred to as the *control samples*. In addition, we generated five random samples of the same number of HA sequences, extracted sequences for the other segments, and reconstructed phylogenies under the same workflow. These random samples are meant to be more consistent with a conventional phylogenetic study.

To visually assess the potential extent of reassortment in the evolutionary history of avian H5Nx genomes, we generated tanglegrams for the HA tree against the trees of every other segment. A tanglegram is a graphical method that re-roots and rotates branches in two trees with matching labels to minimize the discordance in the vertical positions of their labels. Discordance can be quantified by an entanglement statistic that sums the difference in the vertical positions of each label, and normalizes this total such that the statistic ranges from 0 (completely congruent) to 1 (worst case scenario). For example, the tanglegram between trees relating HA and NA segments from a random sample of genomes demonstrates a substantial level of entanglement (0.13; Figure 1A), which implies multiple reassortment events have occurred in the evolutionary history of these sequences. However, these results do not indicate how much discordance can be attributed to inaccuracies in reconstructing the respective phylogenies. To illustrate, Figure 1B displays a tanglegram between the same HA tree and a tree relating sequences simulated along the HA tree adjusted to the sequence length and molecular clock of the segment encoding NA. Since the underlying phylogenies are congruent, this simulations provide an estimate of the expected amount of phylogenetic discordance in the absence of reassortment. Entanglement values were significantly lower for simulated trees (−0.12, 95% confidence interval −0.14, −0.10; Figure 2). Put another way, a tree reconstructed from sequences simulated without reassortment was about 48% less ‘entangled’ than a tree from actual sequences. We also observed significantly lower entanglement on trees generated on random samples (−0.04, 95% CI −0.07, −0.01) relative to control samples. However, entanglement is not a reliable statistic for measuring phylogenetic discordance (De Vienne 2019), and is difficult to interpret in terms of how much reassortment may have occurred between segments.

**Figure 1:**
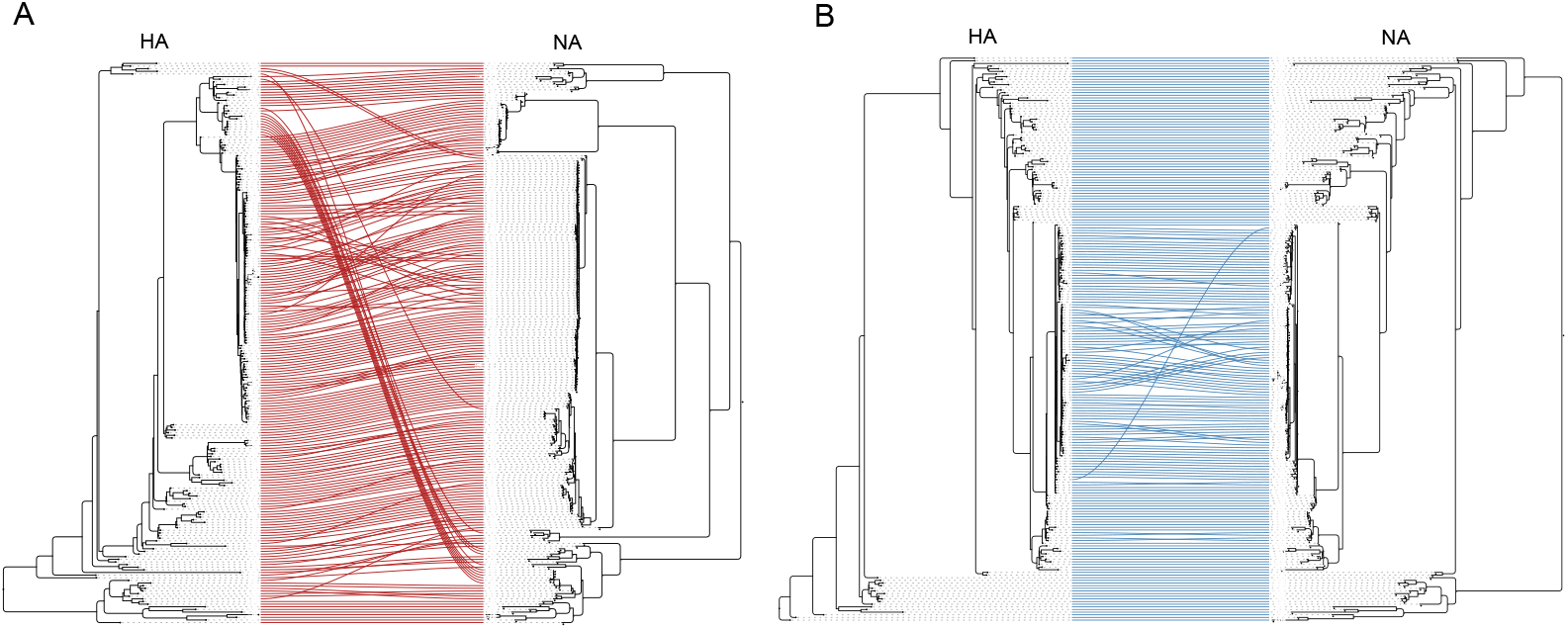
Tanglegrams between a random sample of *n* = 213 influenza A virus H5Nx sequences for segments 4 (HA) and 6 (NA). The NA tree (right side) was reconstructed either from actual segment 6 sequences from the same isolates (A), or generated by simulating sequence evolution along the rescaled HA tree. The entanglement values associated with these pairs of trees were 0.13 (actual) and 0.07 (simulated), respectively. An alternate version using the Python module *baltic* is provided as Supplementary Figure S3.

**Figure 2:**
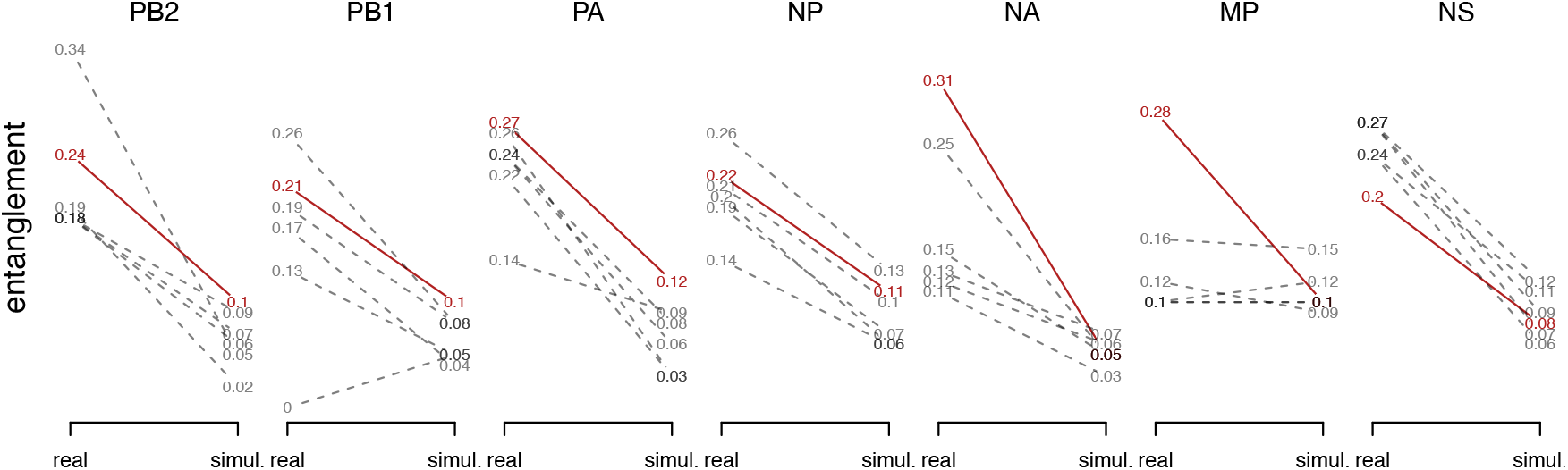
Slopegraphs summarizing entanglement values for real and simulated trees. Solid (red) lines indicate entanglement values obtained for the control (maximum diversity) sample of HA sequences, and dashed lines indicate random samples. ‘Real’ values were obtained for trees reconstructed from actual sequences for the other segments. Simulated (‘simul.’) values were obtained for trees reconstructed from sequences simulated along the rescaled HA tree.

A more robust and informative method for quantifying the discordance between phylogenies is the minimum number of subtree-prune-regraft (SPR) rearrangements required to transform one tree topology into another. This number defines an edit distance between the trees. A useful characteristic of SPR is that this rearrangement is similar to the effect of reassortment on trees Svinti et al. 2013. We calculated the SPR distances between the same pairwise comparisons of trees as we used for entanglement. As expected, the effects of reassortment and phylogenetic reconstruction error were much clearer using SPR counts (Figure 3). SPR counts were significantly lower for simulated trees without reassortment (Poisson regression, *P <* 10^−12^, 95% CI −1.59, −1.45). For random samples of HA sequences, trees reconstructed from other segments simulated without reassortment had about 32.0% (interquartile range, IQR = 17.8% − 36.7%) of SPRs as the real trees, on average. Specifically, the median SPR distance was 98 (IQR 77.5, 103) for real trees and 22 (16, 27.5) for simulated trees (*n* = 35). These proportions were much lower for the control samples (3.8%, range 1.4% − 5.5%), supporting the hypothesis that phylogenetic uncertainty is the main driver of false positives. For these *n* = 7 samples, the median SPR distances were 143 (range 127, 150) for real and 5 (2, 7) for simulated trees. In addition, we obtained consistently higher SPR counts across segments for control samples (mean 140.7, range 127 − 150; Figure 3), which was likely driven by more complete coverage of the underlying evolutionary history of avian H5Nx isolates.

**Figure 3:**
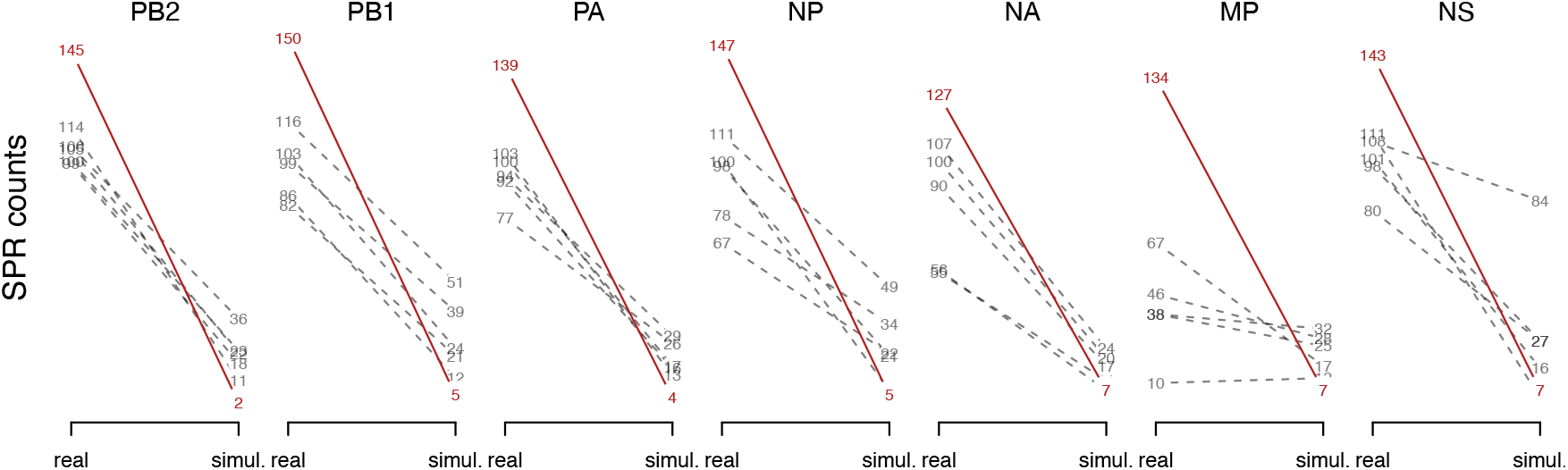
Slopegraphs summarizing subtree-prune regraft distances (minimum SPR counts) between real and simulated trees. Solid (red) lines indicate SPR counts obtained for the control sample of HA sequences, and dashed lines indicate random samples. ‘Real’ counts were obtained for trees reconstructed from actual sequences for other segments. Simulated (‘simul.’) counts were obtained for trees reconstructed from sequences simulated along the rescaled HA tree.

Even though we expect less phylogenetic discordance in the absence of reassortment, the number of false positives is unacceptably high (*∼* 30%) if the sequences have not been down-sampled to maximize evolutionary divergence. Therefore, it would be helpful to be able to distinguish between real and spurious SPR events. We investigated two characteristics of SPR events to determine whether SPRs caused by error in phylogenetic reconstruction were distinguishable from those caused by reassortment. First, we examined the number of terminal nodes (tips) affected by the SPR, *i.e*., subtree size. An unexpected result is that SPRs involving simulated trees tended to have the largest subtrees (Figure 4), despite there being significantly fewer subtrees extracted from these comparisons. This difference was statistically significant for random samples (paired Wilcoxon rank sum test, *P* = 4.16^−6^) and marginally significant for control samples (*P* = 0.051). On the other hand, there was no significant difference between real and simulated trees in the proportion of SPRs involving a single branch (binomial regression, *P* = 0.76).

**Figure 4:**
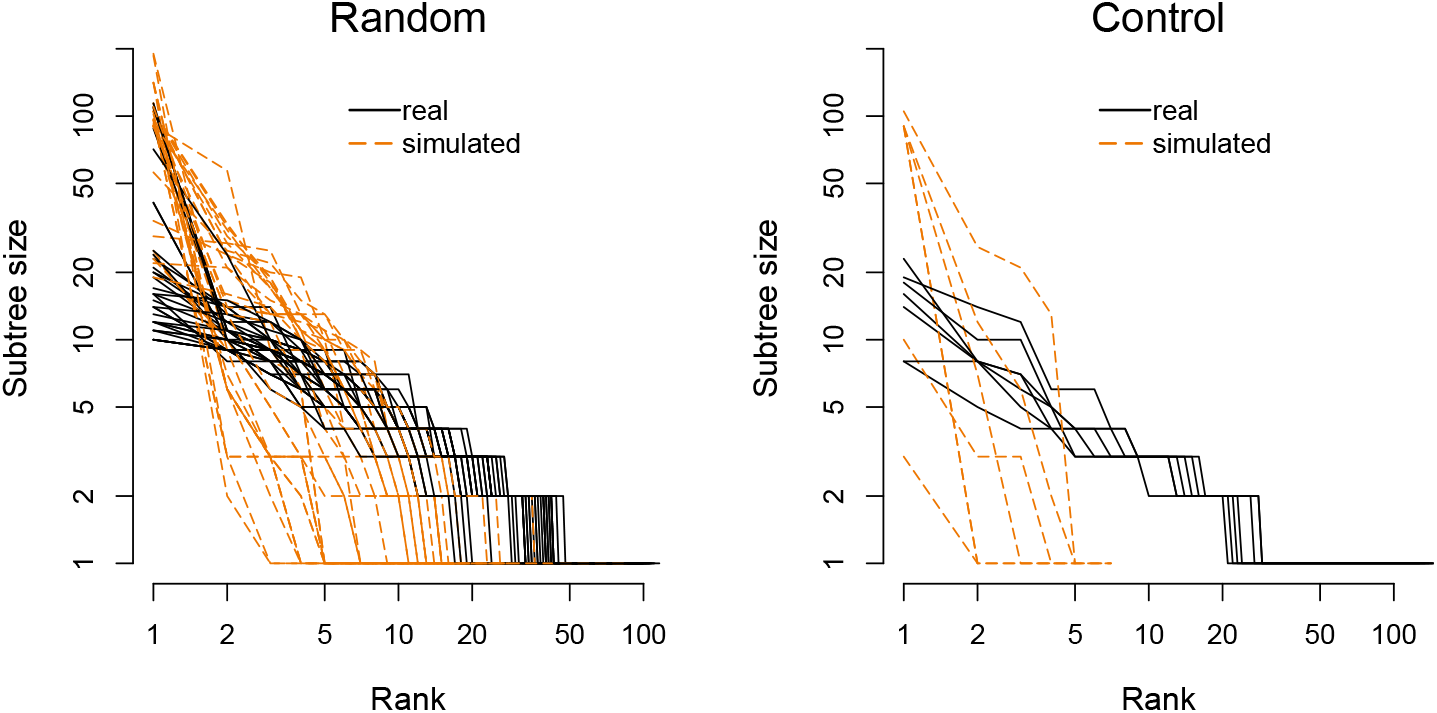
Log-log plots of subtree sizes for all SPR events reconstructed from random and control samples. Each line represents the sequence of subtrees for each segment and replicate (for random samples), sorted in decreasing order of size (number of tips). Solid (black) lines indicate subtrees in trees relating real segments, and dashed (orange) lines indicate simulations. The ranks of the ordered subtrees are plotted along the *x*-axis (*i.e*., largest subtree is rank 1), and subtree sizes are plotted along the *y*-axis.

Next, we quantified the distance between the prune and regraft points for all inferred SPR rearrangements. This task is not straightforward because these points exist on separate trees, so we approximated the path length by locating the smallest subtree containing the same tip labels as the descendants of the sibling node in a second tree, where the sibling node shares the same immediate ancestor as the subtree affected by the SPR. We refer to this path length as the sibling distance for a given SPR. Figure 5 summarizes the distributions of sibling distances between real and simulated trees for random and control samples. Overall, sibling distances tended to be shorter for comparisons to simulated trees, with a median distance of 0.046 for real trees and 0.006 for simulated trees. If we assume that a reassortment event can occur between any two contemporaneous lineages in the population, then a tendency towards shorter sibling distances is more consistent with error in phylogenetic reconstruction. Sibling distances were more variable for comparisons between HA and NA trees (Figure 5), most likely because trees reconstructed from NA sequences were substantially larger than the other segments; conversely, trees relating MP sequences were the shortest. We used a gamma mixed-effects regression model with a log link to test the effect of data type (real or simulated) on sibling distance for random samples, including segment as a random effect. This model obtained a significant negative fixed effect of simulated data on sibling distance for random samples (*P* = 3.1 × 10^−12^, 95% CI −0.53, −0.31). There was significant variation among segments (standard deviation 95% CI 0.28, 0.95). We did not find a significant association between subtree size and sibling distance (*P* = 0.13). Applying the same model to the control samples, we obtained a stronger effect of simulated data (*P* < 10^−12^, 95% CI −2.58, −1.83) and a comparable amount of variation among segments (standard deviation 95% CI 0.29, 1.02). The median sibling distance for control samples was 0.094 for real trees and 0.0078 for simulated trees. These results imply that an effective means of reducing the number of false positives in a reassortment analysis is to down-sample the data to maximize sequence divergence, and to discard SPRs with short sibling distances.

**Figure 5:**
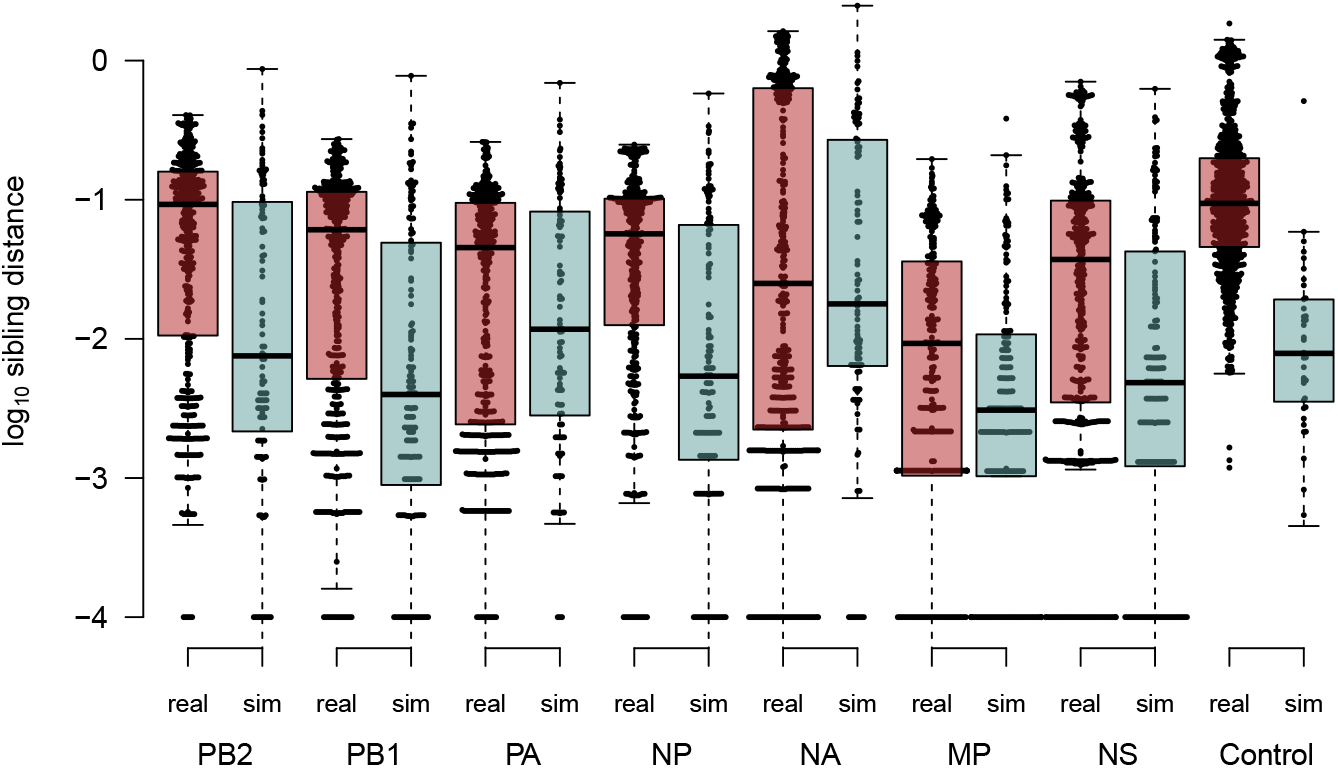
Boxplots showing the distribution of sibling distances (log_10_-transformed) between prune and regraft locations for SPR events across genome segments (PA, PB1, PB2, MP, NA, NP, and NS) for random samples, and pooled across segments for control samples. We added an arbitrary small amount (10^−4^) to zero branch lengths to accommodate the log-transformation.

## Discussion

Reassortment is undoubtedly an important process in the evolution of influenza A viruses in the form of antigenic shift (McHardy and Adams 2009). Several epidemiologically significant strains such as the H1N1 strain associated with the 2009 swine flu pandemic have originated from reassortment combining segments from divergent lineages (Garten et al. 2009; Rambaut et al. 2008). In addition, a wide diversity of influenza A viruses circulate in numerous avian species (Causey and Edwards 2008). This natural host reservoir represents a significant risk for reassortment to generate new genomic combinations with increased pathogenicity and zoonotic potential (Graziosi et al. 2024; Guo et al. 2017; Pulit-Penaloza et al. 2020). For example, Gu et al. (2012) reported an IAV genome isolated from domestic ducks in China in 2012 that appeared to be a reassortment of H11N2, H7N1 and highly pathogenic avian H5Nx lineages. Consequently, there have been many studies using comparative methods to identify reassortment events in the evolutionary history of AIVs. Lewis et al. (2015) reported extensive reassortment from a tanglegram analysis of 211 complete genomes for a variety of HnNn subtypes of AIVs. Similarly, Dugan et al. (2008) observed a “remarkably high rate of genome reassortment” among 167 complete genomes of AIVs, using the difference in log-likelihoods of tree topologies reconstructed from different segments as a measure of discordance when fitting those trees to the same data. Although this method can be used to reject a null hypothesis of no reassortment, it does not identify specific reassortment events. Lu et al. (2014) used ancestral reconstruction of HA or NA subtypes as discrete traits associated with the other segments from about 3,000 AIV genomes to quantify a range of reassortment rates among subtypes and host species. Using a larger dataset of over 40,000 IAV genomes from a variety of host species, Gong et al. (2021) partitioned the trees for the genomic segments into 493-663 monophyletic clades, such that each genome could be represented by the clade assignments of its segments. They reported nearly 2,000 putative reassortments, where a genome was assumed to be a reassortant if its clade assignments could be produced from a combination of two or more other genomes.

Although these studies have employed different methods, they share a common premise on utilizing the phylogenetic discordance among genomic segments as evidence of reassortment. It has long been recognized that error in reconstructing phylogenies will also result in discordant trees in the absence of any reassortment (Svinti et al. 2013). Surprisingly, previous studies of reassortment in AIV genomes have seldom checked for these inevitable false positives. We have found that the estimated prevalence of reassortment events in trees simulated in the absence of any reassortment can reach a substantial fraction (about 20% to 40%) of the number of events estimated from trees derived from actual AIV data. This outcome is sensitive to the amount of divergence in the sequence data. For instance, if the sequences are pre-selected to maximize divergence, making it easier to accurately reconstruct the underlying phylogeny, then the number of SPRs in the absence of reassortment is greatly reduced. This sensitivity to sequence divergence is consistent with error in phylogenetic reconstruction being the primary driver of these false positive SPRs. These results suggest that reports of extensive reassortment among influenza virus segments should be interpreted with some caution. We note that the rapid nucleotide substitution rates of AIV should provide a relative abundance of phylogenetically-informative sites in genomic data within a certain timescale. Identifying reassortment events for other viruses with segmented genomes and slower molecular clocks may be an even greater challenge (Varsani et al. 2018).

In this study, we employed two methods to quantify the extent of reassortment in the evolutionary history of real and simulated data: entanglement and the minimum number of subtree-prune-regraft rearrangements, (*i.e*., the SPR distance). While there are other approaches to quantifying reassortment that have been described in the literature, we submit that entanglement and SPR distances provide a representative selection of methods. Entanglement is generally calculated as a part of generating a tanglegram, a popular method for visualizing phylogenetic discordance (Bansal et al. 2009). It is essentially the total difference in the vertical placement of tip labels. Entanglement can be misleading because the locations of tip labels are not directly proportional to topological congruence. Trees with very different topologies can have a relatively low entanglement, and vice versa (De Vienne 2019; Briand et al. 2020). Our results are consistent with these findings. Specifically, we observed significantly lower entanglement values for trees simulated in the absence of reassortment (Figure 2). However, this pattern was much more clearly resolved for the same data by SPR distances (Figure 3), which is arguably the most direct measure of reassortment that is readily available to us (Svinti et al. 2013; Lin et al. 2024).

A number of programs have been implemented to measure reassortment from phylogenetic discordance (Suzuki 2010). However, only a few of these programs remain available, particularly for inferring the minimum number of SPR rearrangements between trees. For example, SPRSupertrees (Whidden et al. 2014) is a C++ program distributed under the GNU GPLv3 license that calculates the SPR distance, but its computing time because prohibitively long for phylogenetic trees with hundreds of branches or more. In addition, the website from which this program is distributed was not consistently accessible (last attempt October 15, 2025). Espalier (Rasmussen and Guo 2023) uses a heuristic algorithm to compute the SPR distance in a more computationally efficient manner, retaining only strongly supported rearrangements. However, the algorithm employed by Espalier is approximate and can produce distances that are not symmetric, *i.e*., *d*(*A, B*) ≠ *d*(*B, A*).

Lin et al. (2024) recently described several other issues in estimating the number and distribution of reassortment events from discordant segment trees. First, they noted that reassortment events may alter branch lengths without modifying the topology of the trees. Reassortments that involve an unsampled lineage cannot be reconstructed by comparing trees. In addition, comparing discordant trees can be affected by a ‘multiple hits’ problem in which a more recent reassortment event obscures other events that precede it. These issues have the opposite effect compared to phylogenetic uncertainty: they would cause the estimated number of SPR rearrangements to underestimate the actual extent of reassortment. Consequently, measuring the overall rate of re-assortment or reconstructing a number of these events by the large-scale comparative analysis of genomes should be considered an open problem for influenza viruses and other segmented viruses. Fortunately, there has been a resurgence of interest in ancestral recombination graphs (*e*.*g*., Rasmussen and Guo 2023; Barrat-Charlaix et al. 2022; Lewanski et al. 2024) that will likely drive further advances in this area.

## Acknowledgments

We thank David Rasmussen for providing advice and insight into the algorithms employed by Espalier, and Sarah Otto for constructive comments on earlier versions of this manuscript.

## Supplementary Figures

**Figure S1:**
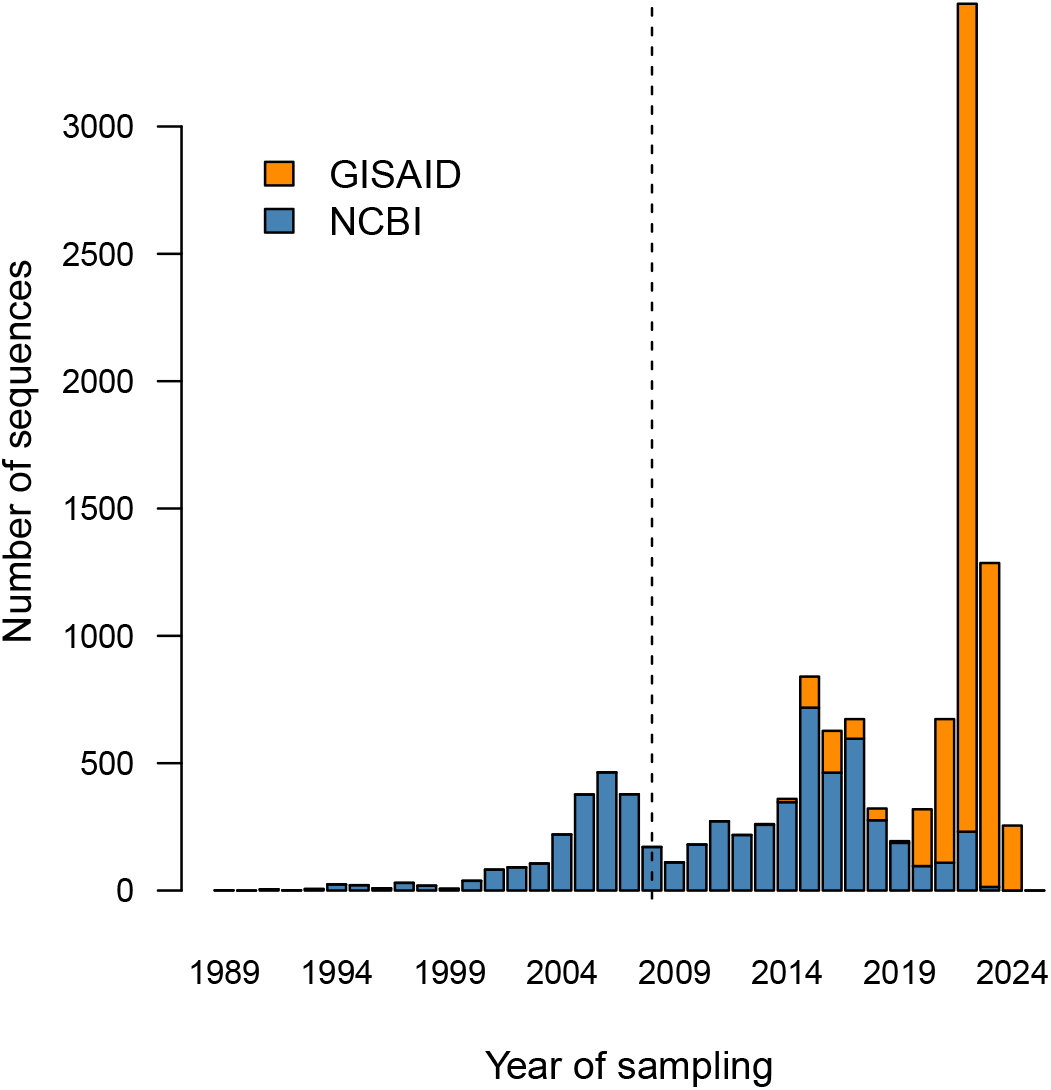
Distributions of sample collection dates for hemagglutinin (HA) sequences from the NCBI Genbank and GISAID databases.

**Figure S2:**
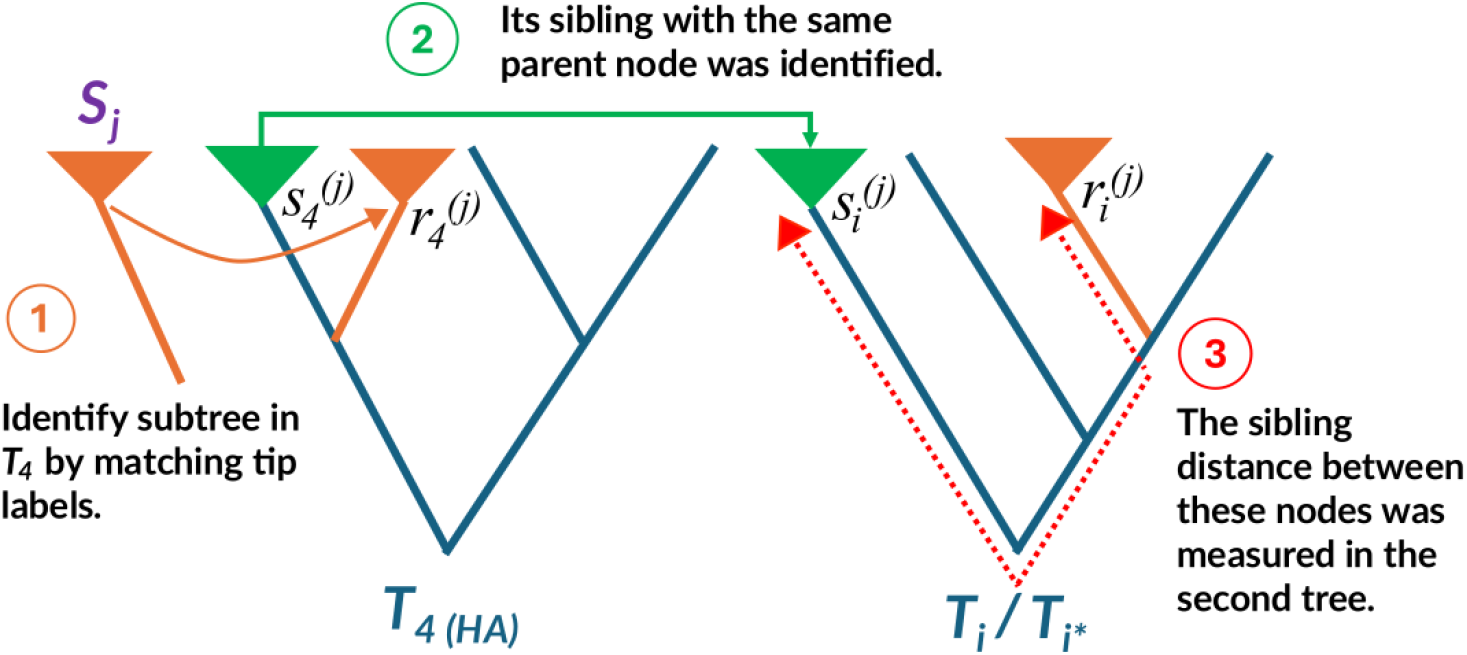
Diagram of the process for calculating the sibling distance of a subtree-prune-regraft (SPR) operation. For each subtree (*S* _*j*_) in the maximum agreement forest, we identify the corresponding subtree in the segment 4 (HA) tree (*T*_4_) by locating the internal node that is ancestral to the smallest set of tip labels matching all labels in *S* _*j*_. Next, we locate the sibling internal node in 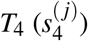 that shares an immediate common ancestor with 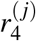. We use the same label matching method to find the analogues of both 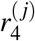 and 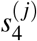 in a second tree relating either real (*T*_*i*_) or sim-ulated (*T*_*i*\*_) sequences of the *i*-th segment (where *i* ≠ 4). Finally, we extract the total path length between these nodes (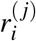 and 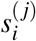) as the sibling distance of the SPR.

**Figure S3:**
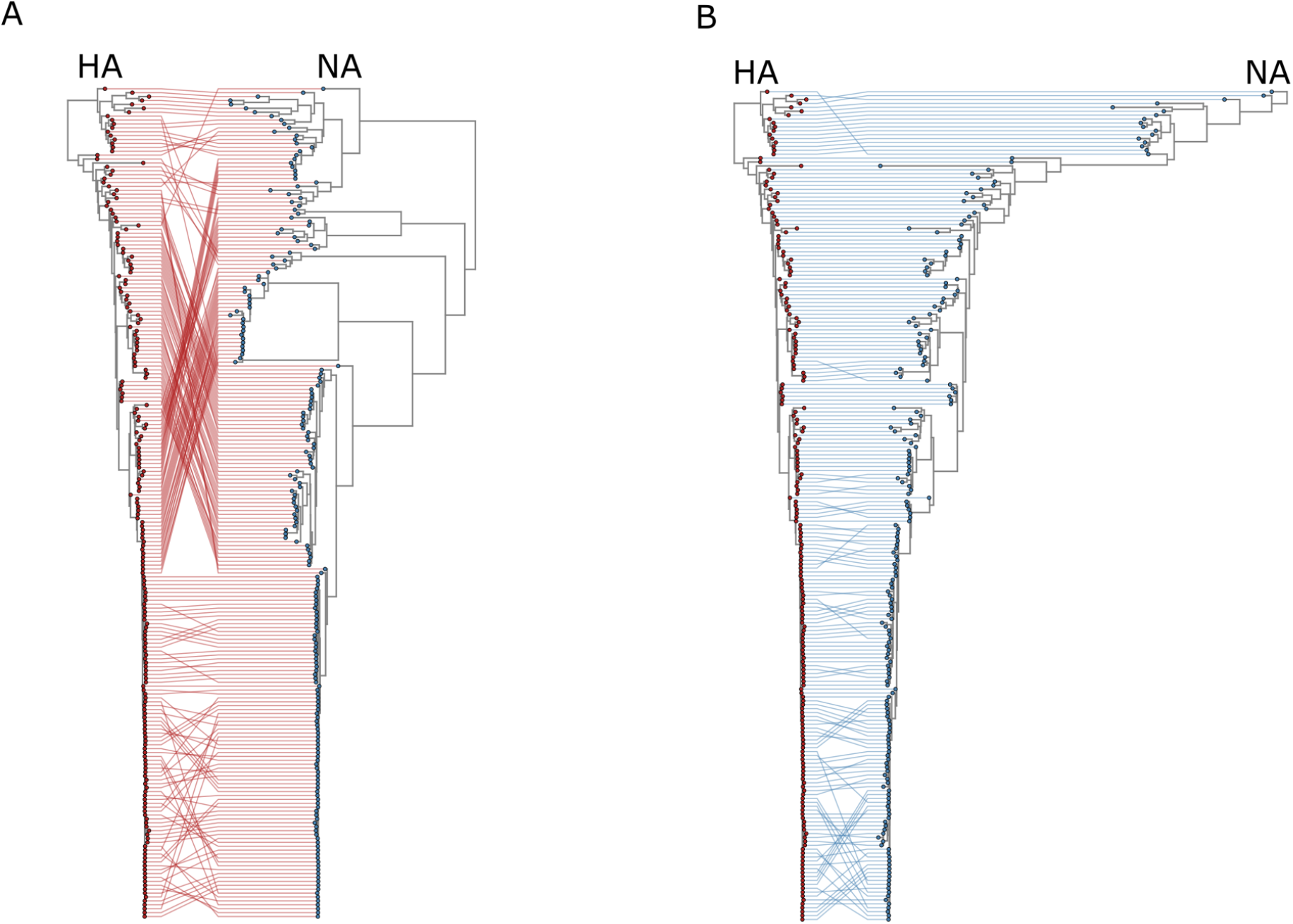
Tanglegram visualizing the correspondence between phylogenetic trees of the HA and NA segments of influenza viruses, generated with the Python module *baltic* (https://github.com/evogytis/baltic). (A) Tanglegram based on real data, showing the relationship and potential reassortment events between matched strains for each segment. (B) Tanglegram comparison of the HA tree real with the simulated scaled NA. This panel shows potential reassortment events that arise in the simulated data.

